# Attomolar fecal cytokine profiling reveals gut immune dynamics and disease states

**DOI:** 10.64898/2026.03.31.714463

**Authors:** Stephanie J. Zhang, Utkarsh Sharma, Yasmeen Senussi, Arvin Dayao, Mia Sveen, Madison Brown, Thanpisit Lomphithak, My Nguyen, Aleigha R. Lawless, Camille A. Briskin, Tatyana Sharova, Genevieve M. Boland, Sonia Cohen, Scott Snapper, Francesca S. Gazzaniga, Lynn Bry, David R. Walt, Travis E. Gibson

**Affiliations:** Pathology Department, Brigham and Women’s Hospital, Boston, MA, USA; Harvard Medical School, Boston, MA, USA; Wyss Institute for Biologically Inspired Engineering at Harvard University, Boston, MA, USA; Broad Institute of MIT and Harvard, Cambridge, MA, USA; Massachusetts Institute of Technology, Cambridge, MA, USA; Division of Gastroenterology, Hepatology, and Nutrition, Boston Children’s Hospital, Boston, MA, USA; Department of Pathology, Massachusetts General Hospital, Boston, MA, USA; Krantz Family Center for Cancer Research, Mass General Brigham Cancer Institute, Boston, MA, USA; Department of Surgery, Mass General Brigham, Boston, Massachusetts, USA; Harvard University, Cambridge, MA, USA; Division of Gastroenterology, Hepatology, and Endoscopy, Brigham & Women’s Hospital, Boston, MA USA

## Abstract

The gut modulates systemic health, influencing immune, neurological, and cardiovascular processes. While fecal sequencing of microbial nucleic acids provides a non-invasive view of microbial composition, sensitive measurement of host-derived signals in stool remains limited. Here we introduce DIGEST (Digital Immunoassay for Gut-Environment Single-molecule Targets), an ultrasensitive digital immunoassay that quantifies proteins in fecal extracts to attomolar levels. In mice, longitudinal profiling during a high-fat diet perturbation revealed coordinated host cytokine responses that occurred within 24 hours, with sustained elevation after diet withdrawal, enabling non-invasive tracking of within-subject immune dynamics. Application of DIGEST to quantify a panel of host inflammatory cytokines in patients with inflammatory bowel disease distinguished active ulcerative colitis from quiescent disease and non-IBD controls (AUC=0.98). In advanced melanoma patients receiving PD-1 blockade, pretreatment fecal IL-23 concentrations discriminated responders from non-responders with an AUC of 0.87. Together, these results establish DIGEST as a generalizable platform for sensitive, non-invasive quantification of host protein activity at the gut interface, with broad applications in basic science discovery, disease surveillance, and therapy response prediction.

## Introduction

The role of the gut in the etiology of diverse disorders has been established over the last two decades, with mounting evidence linking it to diseases far beyond the gastrointestinal tract, including neurological conditions such as Alzheimer’s disease^1,2^ and autism spectrum disorder^3^, autoimmune diseases including rheumatoid arthritis^4^ and multiple sclerosis^5,6^, and gastrointestinal disorders such as inflammatory bowel disease^8^, celiac disease^9^, and colorectal cancer^10^. Despite this growing recognition, current research and diagnostic tools have yet to fully capitalize on the gut’s potential as a window into host physiology. However, studies are often limited to invasive biopsies^11,12^ or peripheral blood analysis, frequently requiring ex vivo stimulation to measure host responses^13^.

Measuring host transcripts in feces is possible with optimized extraction followed by Reverse Transcriptase (RT) quantitative Polymerase Chain Reaction (qPCR), RT-qPCR^14^, or targeted amplification and sequencing approaches^15-18^. However, these studies consistently demonstrate two limitations: (i) given the abundance of microbial transcripts relative to host RNA in fecal samples, host-targeted amplification is required to enrich for low-abundance host signals; and (ii) even with amplification, many host transcripts are not reliably detectable unless the host is undergoing acute infection or active disease.

In contrast, proteins are typically orders of magnitude more abundant than transcripts in human cells^19^, yet remain difficult to quantify in feces as levels typically fall below the detection limits of standard enzyme-linked immunosorbent assays (ELISA)^20^. While ultrasensitive protein assays have yielded remarkable insights in serum^21^, urine^22,23^, and cerebrospinal fluid^24,25^, these technologies have not been adequately extended to fecal samples as a means to more directly interrogate gut responses.

Here we present DIGEST (Digital Immunoassay for Gut-Environment Single-molecule Targets), a digital protein assay developed by adapting ultrasensitive single-molecule detection technologies, including single molecule arrays (SIMOA)^21^, molecular on-bead signal amplification for individual counting (MOSAIC)^26^, and barcoded MOSAIC (b-MOSAIC)^27^, to fecal matrices. Key technical advances include a robust fecal protein extraction workflow and a unified quantitative, matrix-matched calibration framework that replaces separate digital and analog regimes with a continuous, cross-platform quantitative readout. To demonstrate the utility of our assay, we applied DIGEST to a longitudinal mouse study (M1) as well as two human studies (H1 and H2). In M1, we demonstrated DIGEST’s ability to track rapid changes in cytokine levels over many orders of magnitude in a murine model following a High-Fat Diet (HFD) perturbation. In cohort H1, DIGEST differentiated three participant groups: Ulcerative Colitis (UC) patients with active flare (UC+), UC patients without flare at the time of collection (UC−), and healthy controls. Finally, in an exploratory melanoma cohort (H2), pretreatment fecal cytokine levels predicted response to Immune Checkpoint Inhibitor (ICI) therapy with high accuracy. All our analysis and figures can be reproduced from the GitHub repository for this project. https://github.com/gibsonlab/DIGEST

## Results

### DIGEST assay development and analytical performance

To motivate protein-based readouts, we first assessed whether host cytokine transcripts are reliably detectable in stool. Using RT–qPCR for TNF-α in fecal material from conventional healthy mice and a mouse with a *Clostridioides difficile (C. diff)* infection (**Fig. 1a**), most replicates from conventional mice fell below detection threshold (Ct > 35), whereas transcripts from the infected mouse were readily detectable (Ct ∼27), consistent with prior reports that cytokine transcripts in stool are often undetectable outside acute infection or severe inflammation^18^.

**Figure 1:**
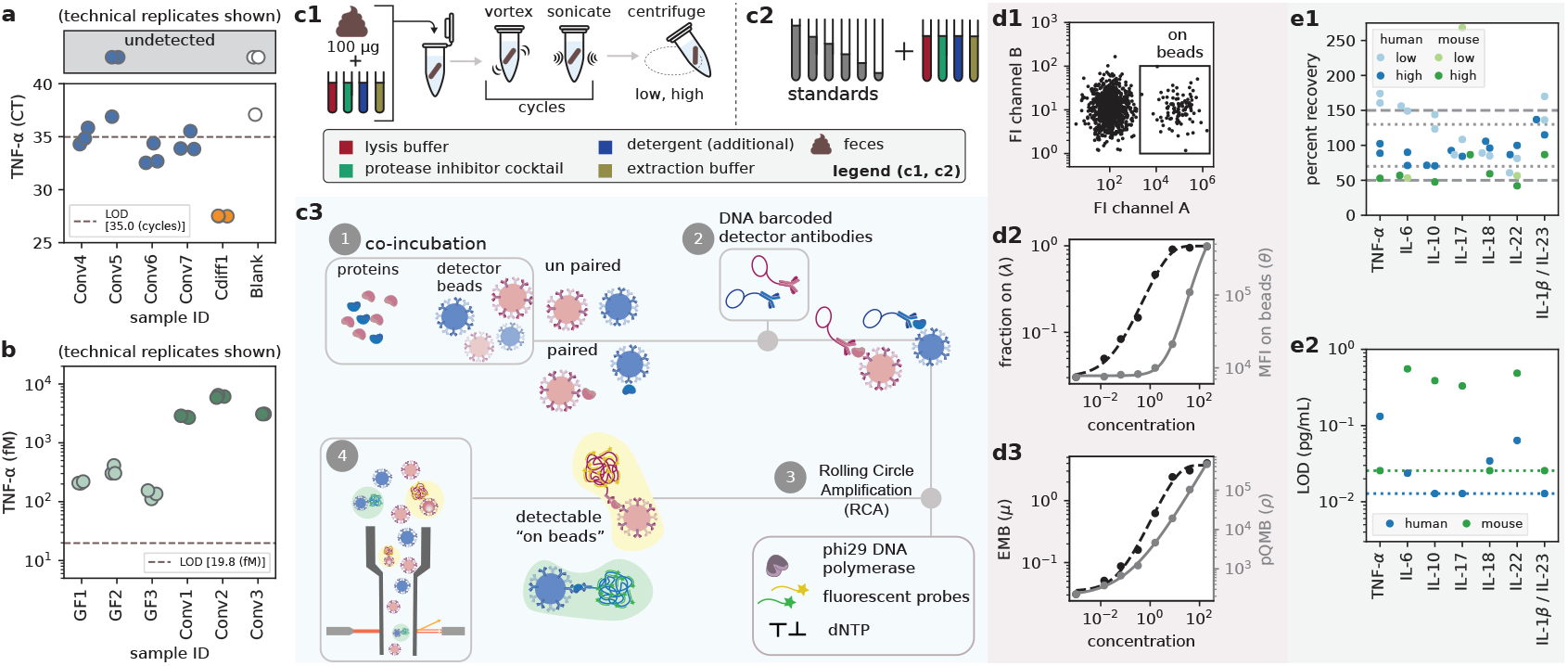
Motivation and overview of DIGEST. **(a)** RT–qPCR cycle threshold (Ct) values for TNFα measured from stool using three technical replicates (all shown) for four conventional mice, one *Clostridioides difficile* (*C. diff*) infected mouse and one blank (no-template control). Note that Ct is inversely proportional to concentration. Replicates in the gray box exceeded the maximum number of cycles without crossing the fluorescence threshold. **(b)** TNFa protein concentrations from stool measured with DIGEST (v0.1) from three germ-free mice and three conventional mice with three technical replicates (all shown). **(c)** DIGEST (v1.0) overview. (c1) Calibration standards are prepared in the same extraction and lysis buffers used for sample processing, rather than in a simple buffer alone, to mirror processed samples. (c2) Fecal samples (as little as 100 μg input) are processed using the same extraction and lysis buffers, followed by vortexing, sonication, and clarification by centrifugation. (c3) Overview of b-MOSAIC technology, (1) beads and proteins are co-incubated (2) DNA barcoded detector antibodies are added (3) Rolling Circle Amplification (RCA) is performed and (4) flow cytometry. **(d)** Flow-cytometry analysis and quantification. (d1) Beads are gated by size and color, with “on” beads identified by probe fluorescence. (d2) From the “on” beads, both the fraction of on beads, *λ*, and the median fluorescent intensity (MFI, *θ*) are recorded. (d3) A unified metric, pQMB (*ρ* = *λ* ⋅ *θ*), is used for calibration across the full dynamic range, shown in comparison to the expected number of molecules per bead (EMB, *μ*) derived from *λ*. **(e)** Spike-and-recovery and analytical sensitivity. (e1) Spikes were performed on a single mouse sample at concentrations of 3.2 pg/mL and 80 pg/mL for each cytokine and for two different human samples with both 1.6 pg/mL and 40 pg/mL. Percent recovery was not plotted when endogenous concentrations exceeded the spike level. (e2) Limits of detection (LOD) for both mouse and human assays were calculated following Clinical and Laboratory Standards Institute (CLSI) recommendations using statistics from blanks and low-concentration standards^7^. Dashed lines indicate the lowest calibration standard, which lower-bounds the LOD.

By contrast, TNF-α protein was readily quantifiable in fecal extracts from both conventional mice (∼10^3^ fM) and germ-free (∼10^2^ fM) using early DIGEST assays (v0.1), well above the run-specific limit of detection (LOD, 19.8 fM, **Fig. 1b**). These data demonstrate that host proteins can be robustly measured in feces even in the absence of corresponding transcripts.

We evaluated three DIGEST variants during development:

- DIGEST (v0.1) – fecal-optimized SIMOA (f-SIMOA)
- DIGEST (v0.5) – fecal-optimized MOSAIC (f-MOSAIC)
- DIGEST (v1.0) – fecal-optimized b-MOSAIC (fb-MOSAIC)

Given autofluorescent particulates in stool that interfere with image-based detection, subsequent development focused on flow cytometry–based platforms.

The final DIGEST v1.0 workflow is shown in **Fig. 1c**. Briefly, fecal inputs as low as 100 *μ*g are extracted and clarified prior to single-molecule analysis (**Fig. 1c1**). Calibration standards are prepared in a matrix-matched format, yielding conservative but representative LOD estimates (**Fig. 1c2**). In DIGEST v1.0, proteins are captured on beads, labeled with DNA-barcoded detector antibodies, and amplified via rolling circle amplification prior to flow cytometric readout (**Fig. 1c3**). Relative to MOSAIC, b-MOSAIC enables greater multiplexing flexibility and reduced nonspecific binding^28^.

Rather than switching between digital and analog calibration regimes, we define a unified metric, pQMB (*ρ* = *λ* ⋅ *θ*), which integrates the fraction of “on” beads (*λ*) and their median fluorescence intensity (MFI or *θ*) to provide a single, continuous calibration across the full dynamic range (**Fig. 1d**). At low concentrations, pQMB coincides with Poisson-derived estimates of molecules per bead (**Fig. 1d3**), while at higher concentrations it maintains log–log linearity without saturation (**Fig. 1d2**). All DIGEST v0.5 and v1.0 measurements in this study use pQMB-based calibration.

Spike-and-recovery (S&R) experiments were performed using one mouse sample and two human samples across seven cytokines (**Fig. 1e1**). Recovery was generally higher in human samples and improved at higher spike concentrations. For human samples, most high-spike recoveries fell between 70– 130%, with low-spike recoveries largely between 50–150%. In mouse samples, high-spike recoveries clustered near ∼50%; for cytokines with endogenous concentrations exceeding the low-spike level, percent recovery was not reported (**see Extended Data Tables**). Recovery was calculated relative to target analyte concentration rather than buffer-only spike controls, providing a more stringent assessment of assay performance^29,30^. Corresponding LODs are shown in **Fig. 1e2**. For six of seven human cytokines, LODs were below 0.1 pg/mL, with several below the lowest calibration standard. Murine assay LODs were higher for four shared cytokines.

### Longitudinal monitoring of murine immune responses to High-Fat Diet perturbation

To demonstrate the utility of DIGEST for non-invasive longitudinal monitoring, we profiled immune dynamics in conventional C57BL/6 mice subjected to a HFD perturbation (**Fig. 2a**). Four singly housed mice were monitored over a 25-day period, with HFD administered from days 6 to 19, enabling within-subject comparison before, during, and after dietary challenge.

**Figure 2:**
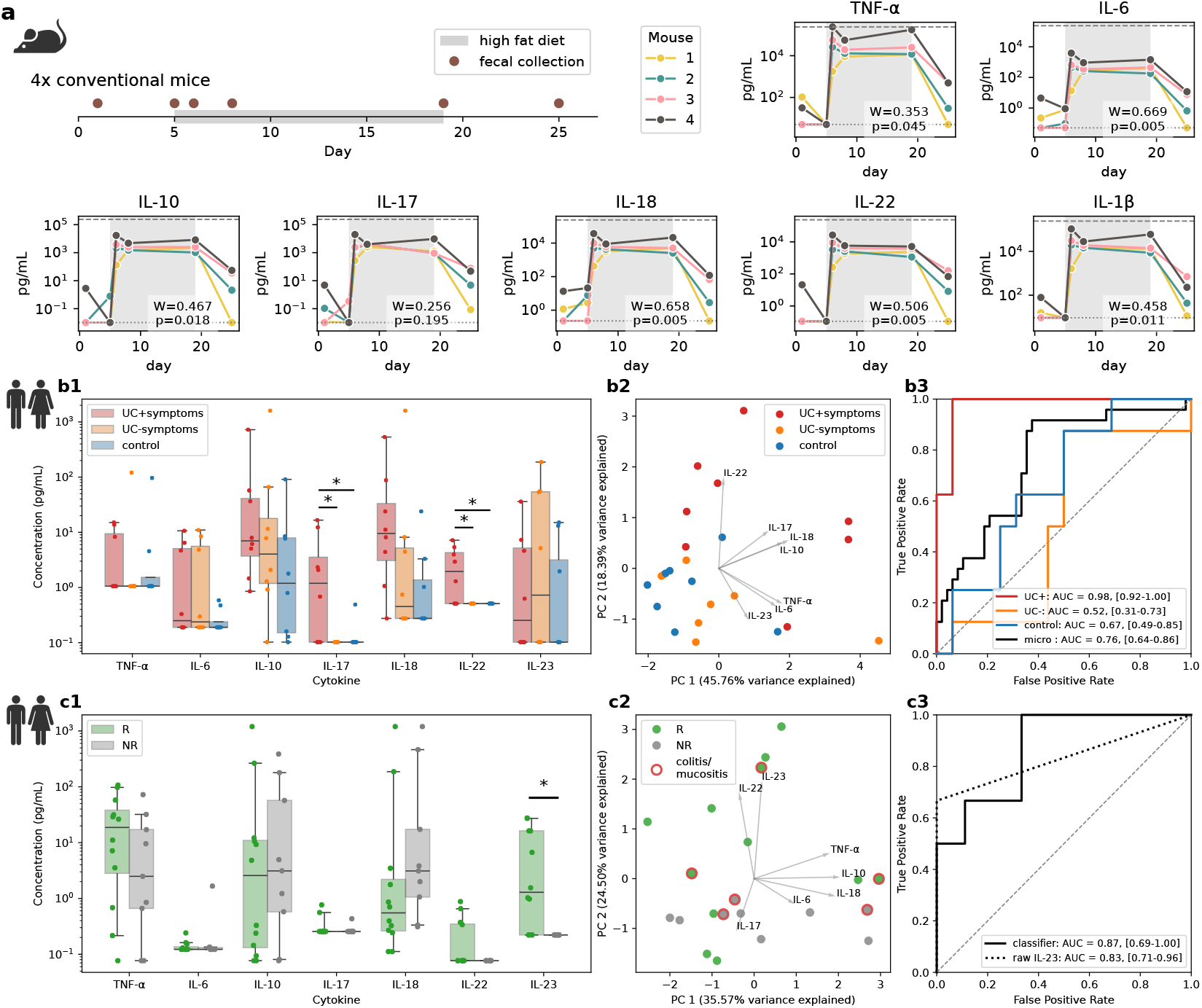
DIGEST enables longitudinal murine immune monitoring and stratifies human disease and treatment-response states from fecal cytokines. **(a) Cohort M1**: Longitudinal murine study design and fecal cytokine dynamics during a high-fat diet (HFD) perturbation in singly housed conventional C57BL/6 mice (4 mice, 25 days; 6 samples per mouse; HFD days administered on 6–19). Shown are the study timeline and fecal cytokine concentrations over time, highlighting a rapid, coordinated cytokine increase within 24 hours of HFD initiation and incomplete return to baseline after diet withdrawal. Rank-order stability across mice was quantified for each cytokine using Kendall’s coefficient of concordance (W) with associated p-value also shown. **(b) Cohort H1**: Human IBD cohort (n=24; 8 UC with active flare [UC^+^], 8 UC without flare [UC^−^], 8 controls). **(b1)** Univariate cytokine differences across groups **(b2)** PCA of the seven-cytokine panel. **(b3)** L1-regularized multiclass logistic regression evaluated by leave-one-out cross-validation (LOOCV), reported as AUC–ROC with 95% confidence intervals (brackets). **(c) Cohort H2**: Advanced melanoma cohort treated with PD-1 blockade (n=21; 12 responders, 9 non-responders). **(c1)** Univariate cytokine comparisons between responders and non-responders with IL-23 elevated in responders. **(c2)** PCA with patients developing ICI colitis/mucositis indicated. **(c3)** LOOCV ROC curves for response prediction using an L1-regularized logistic regression model and a univariate IL-23 classifier, with AUC and 95% bootstrapped confidence intervals shown (brackets). *All p-values shown are Benjamini–Hochberg (BH) corrected. Univariate tests in panels (b1) and (c1) use nonparametric Mann Whitney U test. AUC confidence intervals were estimated by bootstrap resampling under LOOCV. * p < 0*.*05*.

HFD exposure induced a rapid inflammatory response. Within 24 hours of diet initiation, all cytokines in our panel exhibited a synchronized increase spanning four to five orders of magnitude, reflecting an acute, systemic-like shock to the mucosal immune system. This rapid response is consistent with previous reports showing that dietary shifts can alter the gut microbiome and mucosal immune inflammatory state on short timescales^31,32^.

Following HFD withdrawal, cytokine levels subsided but remained above pre-perturbation baselines for a majority of the cytokines and mice, even after six days on normal chow. Despite the modest cohort size, the consistency of these longitudinal responses across animals highlights DIGEST’s sensitivity for resolving longitudinal immune dynamics within individual subjects. This persistence might suggest a degree of “innate immune memory” or what is also referred to as “trained immunity”^33^. Prior work has shown that exposure to saturated fatty acids (specifically palmitate which is found in lard) induces long-lasting transcriptional reprogramming in macrophages via the ceramide pathway^34^. This incomplete recovery also mirrors other work where adipose tissue macrophages have been shown to retain a sensitized, pro-inflammatory phenotype even after dietary normalization in “weight cycling” mouse models^35^.

Visual inspection further suggested stable inter-individual differences in cytokine abundance across mice throughout the study. We quantified this effect using Kendall’s coefficient of concordance (*W*), and observed statistically significant concordance for all cytokines except IL-17, indicating that the relative ordering of cytokine abundances across animals was preserved over time. Notably, these rank orderings remained stable even as absolute cytokine concentrations changed by several orders of magnitude. Such persistent inter-individual immune signatures have been reported in human cytokine profiling studies^36-38^ and here are resolved longitudinally in a controlled murine setting.

### UC flares exhibit an IL-18–IL-17/IL-22–enriched fecal cytokine signature

UC activity reflects fluctuating mucosal immune programs that are difficult to assess non-invasively, and treat-to-target frameworks have emphasized monitoring that incorporates stool-based biomarkers alongside symptoms^39,40^. To test whether fecal cytokine profiling captures UC immune state, we applied our human cytokine panel to fecal samples obtained from an inflammatory bowel disease (IBD) cohort (H1; n=24), comprising UC patients with active flare at collection (UC+), UC patients without flare (UC−), and Controls (n = 8 per group).

Group-wise cytokine profiles are summarized in **Fig. 2b**. IL-17 and IL-22 were significantly elevated in the UC+ samples relative to both UC− and Control (**Fig. 2b1**). In addition, IL-18 and IL-10 exhibited graded increases across groups (Control → UC− → UC+; **Fig. 2b1**). In contrast, IL-23, IL-6, and TNF-α showed comparatively weak separation between groups. These data define a flare-associated fecal cytokine signature characterized by coordinated elevation of IL-18 and IL-17/IL-22, without strong co-variation of fecal IL-23, a cytokine classically associated with Th17 maintenance in chronic IBD pathogenesis^41,42^.

Multivariate analysis reinforced these univariate trends. Principal component analysis separated UC+ samples from UC− and controls along a component co-varying with IL-18, IL-17, IL-22, and IL-10, while IL-23 variance loaded largely orthogonal to this inflammatory cluster (**Fig. 2b2**). Rounding out this analysis, Leave-One-Out Cross Validated (LOOCV) L1-regularized multi-class logistic regression (**Fig. 2b3**) classified UC+ participants with high accuracy (AUC of 0.98, 95% CI: 0.92-1.00), whereas discrimination between controls and UC− was more modest or on par with random guessing.

The observed IL-18–IL-17/IL-22 signature is consistent with an epithelial stress and barrier-associated immune response that, in this cohort, is not tightly coupled to fecal IL-23 levels. IL-18 requires inflammasome- and caspase-dependent processing for activation, positioning it upstream of rapid innate responses to barrier disruption or microbial sensing^43^, and has been reported to track with ulcerative colitis severity^44^. IL-22 promotes epithelial barrier restitution and antimicrobial programs^45^, while IL-17 can arise through IL-23–independent pathways regulating epithelial permeability (e.g. γδ T cell programs)^46^. In addition, IL-18 can expand human ILC3 programs and promote IL-22 production *via* NF-κB, providing a plausible route to an IL-22-high state without requiring strong fecal IL-23 co-variation^47^. Elevated IL-10 in the UC+ group is consistent with prior reports of increased IL-10 expression in active UC mucosa, interpreted as compensatory counter-regulation during inflammation^48^.

All UC+ participants were receiving systemic corticosteroids at the time of collection, with a subset also on anti-TNF therapy (**Human Meta Tables**). This treatment background likely contributes to the muted TNF-α separation observed, as both corticosteroids and anti-TNF therapies suppress canonical TNF-driven inflammatory programs ^49,50^.

### Pretreatment fecal IL-23 predicts ICI response in melanoma

Immune checkpoint inhibitors (ICIs) have transformed cancer therapy, yet response rates remain heterogeneous across indications and patients. In advanced cutaneous melanoma (unresectable stage III/IV), anti–PD-1 monotherapy produces objective response rates on the order of ∼30–40%^51^. Clinical biomarkers such as tumor PD-L1 expression enrich for response in some cancers, including non–small cell lung cancer^52,53^, but predictive markers in melanoma remain limited. Recent studies showing that fecal microbiota transplantation from responders or healthy donors can restore sensitivity to anti–PD-1 therapy in some non-responders and increase responses when administered with anti-PD-1 initiation^54,55^.

Motivated by these observations, we hypothesized that gut-associated immune activity, quantified through fecal cytokine profiles, could provide a non-invasive predictor of ICI response. We analyzed pretreatment fecal samples from patients with advanced melanoma (n=21; 12 responders, 9 non-responders). Cohort-level results are summarized in **Fig. 2c**. Univariate analysis identified IL-23 as the only cytokine significantly different between groups, with higher concentrations in responders (**Fig. 2c1**). TNF-α showed an approximately order-of-magnitude higher median in responders, and IL-18 trended lower, although neither reached statistical significance in this cohort. Multivariate analysis supported these findings. Principal components analysis separated responders from non-responders primarily along PC2 (22.23% variance explained), with IL-23 contributing strongly to this axis (**Fig. 2c2**). Patients who developed ICI-associated colitis or mucositis (blue circles) did not exhibit a distinct pretreatment fecal cytokine signature within this panel.

Using LOOCV L1-regularized logistic regression, fecal cytokines demonstrated discriminatory performance for ICI response with an AUC of 0.87 (95% CI: 0.69–1.00). Notably, a single-analyte model using raw IL-23 concentrations showed comparable classification performance (AUC = 0.83, 95% CI: 0.71–0.96; **Fig. 2c3**). Consistent with our findings, Chen et al.^56^ reported that pretreatment circulating IL-23 levels predicted durable clinical benefit in metastatic melanoma patients receiving nivolumab plus ipilimumab, where IL-23 remained an independent predictor in a “parsimonious” multivariable model (AUC ≈ 0.79)^56^.

IL-23 has been implicated in both pro- and anti-tumor immune processes, and its role in cancer outcomes appears context-dependent^57^. One interpretation of these results is that fecal IL-23 reflects baseline gut immune readiness rather than a direct causal driver of tumor response. In this framework, elevated pretreatment fecal IL-23 serves as a non-invasive readout of a host immune state that may be permissive to effective priming and reinvigoration following checkpoint blockade.

## Discussion

DIGEST (Digital Immunoassay for Gut-Environment Single-molecule Targets) is a fecal-optimized digital immunoassay that enables attomolar-scale quantification of host proteins in stool. Its central technical contributions include: (i) a protein extraction workflow compatible with complex fecal matrices and low input mass; (ii) reagent-matched standard curve construction that more accurately reflects the chemical environment of processed samples; and (iii) a unified quantitative readout (pQMB) that extends dynamic range without requiring separate “digital” and “analog” calibration regimes. These advances address two longstanding challenges in stool protein profiling: matrix-induced distortion of antibody-based measurements and the limited sensitivity of conventional immunoassays.

In a longitudinal murine study, DIGEST revealed synchronized cytokine excursions within 24 hours of HFD perturbation, spanning four to five orders of magnitude, followed by incomplete return to baseline after reversion to normal diet. Beyond cross-sectional differences, DIGEST resolved within-subject immune trajectories over time, capturing both rapid perturbation responses and stable inter-individual immune set points. The persistence of elevated cytokine levels, together with stable inter-individual rank ordering over time, is consistent with durable differences in immune set points and aligns with prior observations of trained or “innate” immune memory induced by metabolic and dietary exposures.

In human samples, fecal cytokine profiling captured clinically relevant immune states in two distinct disease contexts. In ulcerative colitis, active flare was characterized by a dominant IL-18/IL-22/IL-17 signature with comparatively weak IL-23 separation, suggesting that innate epithelial stress and barrier-response programs can be detected non-invasively and may not always align with the canonical IL-23-driven chronic maintenance axis. In an exploratory melanoma cohort treated with PD-1 blockade, pretreatment fecal IL-23 was associated with therapeutic response and showed promising predictive performance, supporting the hypothesis that gut-associated immune activation may reflect a host state permissive to effective checkpoint reinvigoration. These results motivate treating fecal cytokines as compartment-specific immune readouts that complement blood-based biomarkers, offering a non-invasive window into local intestinal immune activity that bridges systemic assays and more spatially resolved modalities. Notably, the divergent behavior of IL-23 across UC flare and melanoma response contexts highlights the state- and compartment-dependence of cytokine relevance, underscoring the importance of biologically matched biomarker measurements rather than universal inflammatory signatures. This distinction may help reconcile conflicting reports on cytokine utility across disease settings and supports a framework in which stool-based immune profiling captures aspects of host–environment interaction that are inaccessible to systemic sampling alone.

This study has limitations and highlights opportunities for further development. First, the human cohorts are relatively small and include heterogeneous treatment backgrounds.

Larger prospective studies will be required to validate predictive performance, refine effect size estimates, and define clinically actionable thresholds. Second, expanding panel breadth and integrating DIGEST with matched microbiome, metabolomic, or other multi-omic measurements should enable more mechanistic interpretation and may further improve predictive modeling. Beyond baseline stratification, DIGEST may be particularly well suited for monitoring dynamic immune responses to interventions such as dietary modulation, microbiome-directed therapies, or immunomodulatory treatment escalation, where repeated invasive sampling is impractical.

In summary, DIGEST provides a generalizable and ultrasensitive approach for non-invasive quantification of host-derived immune signals at the gut interface. By enabling non-invasive longitudinal and quantitative profiling in settings where biopsies or repeated invasive sampling are impractical, DIGEST supports both fundamental discovery and translational biomarker development across inflammatory diseases and cancer immunotherapy, while offering a framework to further elucidate host–microbiome–immune interactions.

## Supporting information

All Tables

## Acknowledgements

SZ was supported by NIH F32AG087642

LB was supported by NIH R01 AI15363, R01 AI179807

GMB/SC would like to acknowledge funding from the Adelson

Medical Research Foundation for support

FSG was supported by NIH 1K22CA258960-0, KFCCR Spark

Award, Cancer Research Institute CLIP Award, and the

Victoria’s Secret Global Fund for Women’s Cancers Career

Development Award in Partnership with Pelatonia and AACR

SBS was supported by NIH P30DK034854

DW was supported by NIH 1R01EB032826

TEG was supported by NIH R35GM143056, NSF 2025515.

All reagents and core facility costs were purchased using MGB funds managed by TEG. Gnotobiotic mouse studies were supported by the Harvard Digestive Diseases Center grant P30 DK034854 and a capital grant from the Massachusetts Life Sciences Center.

## Author contributions

Following the CRediT taxonomy standard https://credit.niso.org

### All Categories

Conceptualization, Data curation, Formal analysis, Funding acquisition, Investigation, Methodology, Project administration, Resources, Software, Supervision, Validation, Visualization, Writing – original draft, Writing – review & editing

- SJZ - Conceptualization, Data curation, Formal analysis, Funding acquisition, Investigation, Methodology, Software, Supervision, Validation, Visualization, Writing – original draft, Writing – review & editing
- US - Conceptualization, Data curation, Formal analysis, Investigation, Software, Visualization, Writing – original draft, Writing – review & editing
- YS - Investigation
- AD - Investigation
- MS - Data curation
- MB - Data curation
- TL - Investigation
- MN - Investigation
- ARL - Investigation, data curation
- CAB - Investigation, data curation
- TS - Investigation, data curation
- GMB - Supervision, data curation, resources
- SC - Supervision, data curation, resources
- SS - Supervision, Data curation, Resources, Writing – review & editing
- FG - Conceptualization, Data curation, Investigation, Writing – original draft, Resources, Writing – review & editing
- LB - Supervision, Data curation, Methodology, Resources, Writing – review & editing
- DRW - Conceptualization, Funding acquisition, Methodology, Resources, Writing – review & editing
- TEG - Conceptualization, Data curation, Formal analysis, Funding acquisition, Investigation, Methodology, Project administration, Resources, Software, Supervision, Validation, Visualization, Writing – original draft, Writing – review & editing

## Competing interest statement

DRW is a founder and equity holder of Quanterix Corporation. His interests were reviewed and are managed by Brigham and Women’s Hospital and Mass General Brigham in accordance with their conflict-of-interest policies. In Addition, DRW has a patent for WO2023059731A3 and licensed to Quanterix Corporation. DRW and SJZ have a patent for Cross-Reactivity-Free Single Molecule Assays for Ultrasensitive Detection of Biomolecules with Improved Multiplexing Capability (PCT/US2025/051031) pending to none. GMB has sponsored research agreements through her institution with: Olink Proteomics, Teiko Bio, InterVenn Biosciences, Palleon Pharmaceuticals, Astellas, AstraZeneca. She served on advisory boards for: Iovance, Merck, Moderna, Nektar Therapeutics, Novartis, Replimune, and Ankyra Therapeutics. She consults for: Merck, InterVenn Biosciences, Iovance, and Ankyra Therapeutics. She holds stock options in Ankyra Therapeutics. SBS declares the following interests: Scientific advisory board participation for Pfizer, Trex Bio, Zag Bio, Vendata, Foli Bio; Orikine; Slate Bio, and Ignite Biomedical. SJZ, US, DRW, and TEG report a patent for (64/016,303) pending.

## Code and Data Availability

All figures are reproducible with python notebooks that can be found here: https://github.com/gibsonlab/DIGEST. Raw data used to generate all standard curves is also contained within the GitHub repo.

## Materials and Methods

### Materials and Reagents

All affinity reagents, recombinant proteins, and DNA oligos used in this work are listed in separate tabs within the **Reagent Tables** excel file. Buffers and paramagnetic beads were purchased from Quanterix Corporation and Bangs Laboratories. Custom DNA oligos were purchased from Integrated DNA Technologies and MilliporeSigma.

### Animal Cohorts

All animal studies were conducted under MGB IACUC protocol 2020A011295. Three different groups of mice were used in this study for the (1) RT-qPCR measurements in Figure 1a, (2) DIGEST measurements in Figure 1b, and (3) DIGEST measurements with the diet perturbation model in Figure 2a. For the RT-qPCR measurements fecal samples were collected from five male Swiss Webster mice (four healthy and one with an active *C. diff* infection). For the preliminary DIGEST measurements in Figure 1b samples were collected from six male C57BL/6 mice (three conventional and three germ free). For the longitudinal study four male C57BL/6 mice were singly housed in OptiMice containment cages (Animal Care Systems, Centennial, CO) for the duration of the experiment. The High Fat Diet (HFD) was Teklad (Inotiv) TD.06414 which is approximately 60% of calories from fat (34% of diet by weight) with an energy density of 5.1 kcal/g. The primary source of fat is lard. The normal chow used in this study was Autoclavable Mouse Breeder Diet 5021 from LabDiet. All mice in the study were between 6 and 9 weeks old.

### Human Subjects

All human samples were de-identified, and experiments were conducted under Institutional Review Board (IRB) approval from Mass General Brigham.

Patients with ulcerative colitis and non-inflammatory control subjects were recruited at Boston Children’s Hospital through the Pediatric Gastrointestinal Disease Biospecimen Repository and Data Registry (IRB: P00000529). Demographic and clinical data, including sex, age, body mass index (BMI), fecal calprotectin, C-reactive protein (CRP), Pediatric Ulcerative Colitis Activity Index (PUCAI) score when applicable, and medication use, were collected prior to and at the time of sample collection. Stool samples were collected prospectively in provided containers, stored in home freezers within 1 hour of collection until transport to the hospital, and then stored at -80 °C until further processing.

Fecal samples from advanced melanoma patients treated at Massachusetts General Hospital were collected under a Dana-Farber/Harvard Cancer Center IRB-approved protocol (DFHCC 11-181) and de-identified prior to analysis.

### Fecal RT-qPCR measurements

Samples were collected, flash frozen in liquid nitrogen, and then immediately stored at −80 °C. Two fecal pellets/sample were extracted with the King Fisher Flex instrument using the MagMax Microbiome Ultra Kit. DNA was removed from samples with the Zymo Research RNA Clean and Concentrator-5. cDNA was generated with the Origene First Strand cDNA Synthesis Kit. RT-qPCR performed in triplicate with the PowerUp SYBR Master Mix using the comparative Ct method. TNFalpha mouse primers (used by Origene, ordered from IDT, F-GGT GCC TAT GTC TCA GCC TCT T; R-GCC ATA GAA CTG ATG AGA GGG AG). GAPDH was included as a control and was ordered from IDT as well: primers (F-ACC ACA GTC CAT GCC ATC AC, R-TCC ACC ACC CTG TTG CTG TA).

### Sample preparation for DIGEST measurements

Fecal samples were first weighed in milligrams and transferred into Protein LoBind tubes (Eppendorf). To each 10 mg of fecal material, 100 µL of extraction buffer (0.01 M 1× PBS, pH 7.4, containing 0.5% Tween-20 and 0.05% sodium azide) and 100 µL of assay buffer (1% Triton X-100, 1× protease inhibitor cocktail, and 1× EDTA in Quanterix sample diluent) were added to facilitate protein solubilization and stabilization. To ensure thorough disruption of the fecal matrix, samples were homogenized by vortex mixing for 10 ∼15 min, followed by 5 min of sonication. This process was repeated three times to maximize analyte extraction. The suspensions were then subjected to centrifugation at 10,000 × g for 10 min at 4 °C, after which the supernatant was carefully transferred to fresh Protein LoBind tubes. A second centrifugation step was performed at 10,000 × g for 10 min in a microcentrifuge to further clarify the samples. The final supernatants were collected in fresh Protein LoBind tubes for MOSAIC analysis, with any remaining volume stored at −80 °C for future use. Extraction and processing conditions were optimized based on previous literature^58,59^. Calibrator standards were prepared in the same extraction and assay buffer mixture used for fecal samples to ensure matrix-matched calibration.

### Preparation of capture and labeling reagents

Capture antibodies were buffer exchanged with Bead Conjugation Buffer (Quanterix) using a 50K Amicon Ultra-0.5 mL centrifugal filter (MilliporeSigma). After adding Bead Conjugation Buffer to antibody solution in the filter up to 500 µL, buffer exchange was carried out by centrifuging three times at 14,000xg for five minutes, with addition of 450 µL Bead Conjugation Buffer between centrifugation cycles. The buffer-exchanged antibody was recovered by inverting the filter into a new tube, centrifuging at 1000xg for two minutes, rinsing the filter with 50 µL Bead Conjugation Buffer, and centrifuging one more time at 1000xg for two minutes. The concentration of the buffer-exchanged antibody was then measured using a NanoDrop spectrophotometer. For each bead type, beads were washed three times with 300 µL Bead Wash Buffer (Quanterix) and two times with 300 µL Bead Conjugation Buffer (Quanterix) before resuspending in cold Bead Conjugation Buffer. Bead number and conjugation conditions for each analyte are shown in **Reagent Tables** excel file (Table R3 tab). A 1 mg vial of 1-ethyl-3-(3-dimethylaminopropyl) carbodiimide hydrochloride (EDC) (Thermo Fisher Scientific) was dissolved in 100 µL cold Bead Conjugation Buffer, and the desired volume was added to the beads. The beads were shaken for 30 minutes at either room temperature or 4°C. After EDC activation of the carboxyl groups on the beads, the beads were washed once with 300 µL cold Bead Conjugation Buffer before resuspension in the buffer-exchanged antibody solution. Antibody conjugation was carried out by shaking the beads for two hours at either room temperature or 4°C, followed by washing twice with 300 µL Bead Wash Buffer. The antibody-coupled beads were then blocked for thirty minutes at room temperature with shaking in 300 µL Bead Blocking Buffer (Quanterix). After washing once each with 300 µL Bead Wash Buffer and Bead Diluent (Quanterix), the beads were resuspended in Bead Diluent, counted with a Beckman Coulter Z1 Particle Counter, and stored at 4°C.

Unless otherwise indicated by the **BAF-** prefix (pre-biotinylated), all detector antibodies were purchased in unconjugated form and subsequently biotinylated in-house. The detector antibody was biotinylated by reconstituting to 1 mg/mL in Biotinylation Reaction Buffer (Quanterix) and adding a 40-fold molar excess of NHS-PEG4-Biotin (Thermo Fisher Scientific) freshly dissolved in water. The biotinylation reaction was carried out for thirty minutes at room temperature, followed by purification of the biotinylated antibody with a 50K Amicon Ultra-0.5 mL centrifugal filter. Five centrifugation cycles of 14,000xg for five minutes with addition of 450 µL Biotinylation Reaction Buffer between cycles were performed, with subsequent recovery of the purified antibody via inversion of the filter into a new tube and centrifuging at 1000xg for two minutes. The filter was then rinsed with 50 µL Biotinylation Reaction Buffer before centrifuging one more time at 1000xg for two minutes and quantifying the antibody concentration with a NanoDrop spectrophotometer.

### Preparation of streptavidin-DNA conjugate

A 5’ azide-modified primer was annealed to the DNA template for RCA by heating a solution of 33.8 µM primer and 40.6 µM template in NEBNext Quick Ligation Buffer (New England Biolabs) at 95°C for two minutes and allowing to cool to room temperature over 90 minutes. Ligation was then performed with addition of T4 DNA ligase and incubation at room temperature for three hours. The ligation reaction was then heated at 65°C for 10 minutes to inactivate the ligase, with subsequent cooling to room temperature. The ligation reaction was buffer exchanged into phosphate buffered saline (PBS) with 1 mM EDTA using a 7K MWCO Zeba spin desalting column (Thermo Fisher Scientific). For conjugation, streptavidin (Biolegend 280302) was buffer exchanged into PBS using a 10K Amicon Ultra-0.5 mL centrifugal filter, incubated with a 20-fold molar excess of dibenzocyclooctyne-PEG4-N-hydroxysuccinimidyl ester (DBCO-PEG4-NHS, MilliporeSigma) for 30 minutes at room temperature, and purified with a 10K Amicon Ultra-0.5 mL centrifugal filter in PBS with 1 mM EDTA. A two-fold molar excess of the ligated primer-template was then added to the DBCO-modified streptavidin and incubated overnight at 4°C. The conjugate was stored in aliquots at -80°C in PBS with 5 mM EDTA, 0.1% BSA, and 0.02% sodium azide.

### MOSAIC assays

MOSAIC assays were performed in a 96-well plate (Greiner Bio-One, 655096), with antibody-coated beads and detector antibodies diluted to the desired concentrations in Homebrew Sample Diluent (Quanterix). Assay conditions for each analyte are listed in **Reagent Tables** excel file (Table R3 tab). Sample volumes of 100 µL were used, with 10 µL of antibody-coated beads. For two-step assays, 10 µL detector antibody was added to each sample. The plate was sealed and shaken for one hour for target capture, followed by washing with System Wash Buffer 1 (Quanterix) using a BioTek 405 TS Microplate Washer. For three-step assays, 100 µL detector antibody was added to the beads after target capture and washing steps, followed by a 10-minute incubation and additional washing. The immunocomplex sandwiches were then labeled with streptavidin-DNA by adding 100 µL of the conjugate diluted in Sample Diluent with 5 mM EDTA to the beads and shaking for the desired time. The samples were then washed with System Wash Buffer 1 for eight cycles, transferred to a new 96-well plate (Greiner Bio-One, 655901), and washed an additional time with 200 µL System Wash Buffer 1 before being resuspended in 60 µL of the RCA reaction mixture. The RCA mixture consisted of 0.5 mM deoxynucleotide mix (New England Biolabs), 0.33 U/uL phi29 DNA polymerase, 0.2 mg/mL bovine serum albumin (BSA, New England Biolabs), 1 nM fluorescently labeled DNA probe (Integrated DNA Technologies), and 0.1% Tween-20 in 50 mM Tris-HCl (pH 7.5), 10 mM (NH_4_)_2_SO_4_, and 10 mM MgCl_2_. ATTO-647N and ATTO-565 labeled DNA probes were used for single target and multiplex MOSAIC assays, respectively. Upon addition of the RCA mixture to each sample, the plate was shaken for one hour at 37°C, followed by addition of 150 µL PBS with 0.1% Tween-20 and 5 mM EDTA to stop the reaction. Samples were washed one time with 200 µL of the same PBS-Tween-EDTA buffer and resuspended in 100 µL of the buffer with added 0.1% BSA. Samples were measured using a CytoFlex LX flow cytometer (Beckman Coulter) equipped with three lasers, in either tube or plate sampling mode. Bleach and buffer wells were included between different samples to minimize potential sample carryover. Multiplex MOSAIC assays were carried out following the same protocol as for the single-plex MOSAIC assays, with different fluorescent dye-encoded beads combined in the same sample.

Flow cytometry data were first analyzed with FlowJoTM Software (Becton, Dickinson and Company); beads were identified using gates on forward scatter, side scatter, and bead fluorescence. Single beads were additionally gated using forward scatter. The probe fluorescence intensities for each bead population were analyzed using FlowJoTM Software.

### Calibrator standards preparation with matrix-adapted reagents

To minimize matrix-dependent bias and ensure consistency between calibration standards and biological samples, calibration curves were generated using the same extraction and lysis buffers used for sample processing, rather than a separately optimized buffer for recombinant protein standards. Specifically, recombinant analytes were diluted into this same extraction and lysis buffer composition (including 0.01 M 1× PBS, pH 7.4, 0.5% Tween, 0.05% sodium azide, and a lysis mixture containing 1% Triton X-100, 1× protease inhibitor, and 1× EDTA in Quanterix sample diluent) to generate the calibration series. The Quanterix sample diluent contains a proprietary formulation that includes buffering salts (disodium phosphate, monopotassium phosphate, sodium chloride, and potassium chloride), bovine serum albumin, Tween, EDTA, ProClin 300 preservative, and interference-blocking reagents, with exact concentrations not disclosed.

Briefly, biological samples (e.g., fecal samples) were processed using this defined extraction and lysis workflow. Recombinant target analytes were then diluted directly into the same buffer system, rather than being prepared in an idealized diluent. This approach ensures that both standards and biological samples are exposed to closely matched buffer conditions, including detergents, salts, preservatives, and protease inhibitors that may influence protein stability, binding interactions, and assay signal.

### Expected molecules per bead (EMB) under Poisson assumption

The probability mass function for a Poisson distribution parametrized by mean parameter *μ* with *k* molecules per bead is defined as

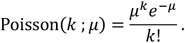

The expected value of the distribution then is simply the mean, 𝔼(Poisson(⋅ ; *μ*)) = *μ*. Therefore, the probability of a bead having zero molecules (“off”) is then Poisson(0 ; *μ*) =e^−u^. If we define the probability that a bead is “on” as *λ* (equivalent to fraction of beads observed to be on), then

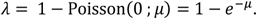

Therefore, if you have measured the fraction of on beads (*λ*) then the expected number of molecules per bead is given by

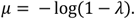

### Standard curve fitting

All standard curves in this study used the five-parameter logistic regression (5PL) model defined as

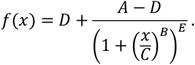

We fit the model using weighted least squares optimizing the following loss function

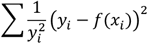

where *x*_*i*_ and *y_i_* are the known concentration and measured signal, respectively.

### Limit of Detection Calculations

We determined the LOD following the Clinical and Laboratory Standards Institute (CLSI) recommendations^7^.

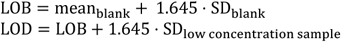

### Spike and Recovery

For spike and recovery the Percent Recovery (PR) in this study is defined as

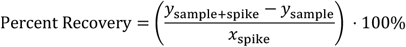

where the measured values are denoted with a *y* and are the measured concentration of the sample with spike-in,*y*_sample+spike_, as well as the measured sample without the spike, *y*_sample_. The target concentration of the spike-in (not a measured value) is denoted as *x*_spike_. When *x*_spike_ was greater than *y*_sample_ we did not report the percent recovery in **Fig. 1**, but the calculated value is still presented in the raw data tables.

We mentioned in the main text that others don’t follow this definition. So for instance, with NULISA they report^30^

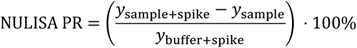

Where *y*_buffer+spike_ is what they measure from spiking into buffer (using the same assay they are trying to validate). Biases in your assay can be recapitulated in the buffer measurement.

### Kendall’s coefficient of concordance (*W*) and permutation test for longitudinal mouse study

We quantified concordance of mouse-specific ordering across repeated timepoints using Kendall’s coefficient of concordance (*W*). For each analyte, we first constructed a timepoint-by-mouse matrix of measurements and then converted measurements to within-timepoint ranks. Within each timepoint *t*, values across mice were ranked using average ranks for ties (midranks).

Let *m* be the number of timepoints and *n* the number of mice, and let *r*_*ti*_ denote the rank assigned to mouse *i* at timepoint *t*. For each mouse *i*, we computed the rank-sum across timepoints,

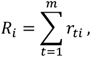

and the expected rank-sum under no across-time concordance,

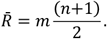

We then calculated

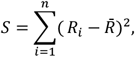

and Kendall’s coefficient of concordance,

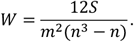

*W* ranges from 0 (no agreement in the mouse ordering across timepoints) to 1 (perfectly consistent ordering across timepoints).

To test whether the observed concordance exceeded that expected by chance, we used a label-shuffling permutation test on the rank matrix. Under the null hypothesis of no consistent ordering across timepoints, mouse identities are exchangeable within each timepoint. We therefore generated *B* permuted datasets by independently permuting the mouse labels (i.e., permuting columns) separately at each timepoint, recomputing *W* for each permutation to form the null distribution 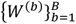. The one-sided p-value was estimated as

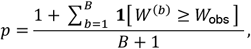

where *W_obs_* is the observed concordance. We used *B* =20,000 permutations. The permutation based p-value was then Benjamini–Hochberg (BH) corrected.

### Univariate Statistical Tests for IBD and ICI cohorts

For the ICI samples responders and non-responders were compared for each cytokine using the Mann-Whitney U test, followed by BH correction. For the IBD samples univariate statistical analysis was performed in a hierarchical fashion. To determine if there were any differences between the three groups (UC+, UC−, control) we first performed a Kruskal-Wallis H test, followed by BH correction. Then for those cytokines that did have significant differences we performed Mann-Whitney U test between all pairs within the three groups for each cytokine. BH correction was then applied to all the Mann-Whitney U test p-values that had been calculated in the previous step.

### Logistic Regression Classifier

All classifiers were L1 regularized logistic regression models trained using nested Leave-One-Out-Crossfold-Validation (LOOCV) on balanced accuracy. Inner folds used for hyper-parameter tuning used three splits (k=3). Concentrations were first log transformed and then subsequently standardized to a normal distribution (mean zero and variance one) using only the training data. Learned transformations were then only applied to test data during testing.

Pseudo code using numpy and scikit-learn libraries including pipeline for the classifier is as follows:

~~~
# X are the concentrations
# y are the class labels
X=np.log(X)
# use pipeline for data standardization and model
specification
pipeline = Pipeline([(‘scaler’,
    StandardScaler()),(‘classifier’,
    LogisticRegression(penalty=‘l1’,
    solver=‘liblinear’))])
#set up inner loop folds
cv=StratifiedKFold(n_splits=3, shuffle=True)
#parameter grid for the inner loop hyperparameter
search
param_grid = {‘classifier C’: np.logspace(-1, 1, 20)}
# initialize LeaveOneOut
loo = LeaveOneOut()
all_probabilities = []
all_true_labels = []
# iterate through each outer fold
for train_i, test_i in loo.split(X):
   X_train, X_test = X[train_i], X[test_i]
   y_train, y_test = y[train_i], y[test_i]
# hyper-parameter search with inner folds
inner_grid_search = GridSearchCV(
   estimator=pipeline,
   param_grid=param_grid,
   cv=cv,
   scoring=‘balanced_accuracy’,
   n_jobs=-1
  )
# fit training data
inner_grid_search.fit(X_train, y_train)
# extract prediction for test data
probabilities =
inner_grid_search.predict_proba(X_test)
# construct loo predictions
all_probabilities.append(probabilities[0])
all_true_labels.append(y_test[0])
~~~

### AUC and Bootstrap confidence intervals

Model performance was assessed by the area under the receiver operating characteristic curve (AUC) using pooled out-of-sample predicted probabilities from the outer cross-validation loop. Confidence intervals for AUC were estimated by nonparametric bootstrap resampling of patient-level outcome– prediction pairs. In each of 10,000 bootstrap replicates, patients were sampled with replacement and the AUC was recalculated. Bootstrap resampling was performed in a stratified manner by sampling outcome-positive and outcome-negative patients separately with replacement while preserving the original class counts. The reported point estimate is the AUC on the full pooled out-of-sample predictions, and the 95% confidence interval was defined by the 2.5th and 97.5th percentiles of the bootstrap distribution.

## Supplementary Figures

**Supplemental Figure S1:**
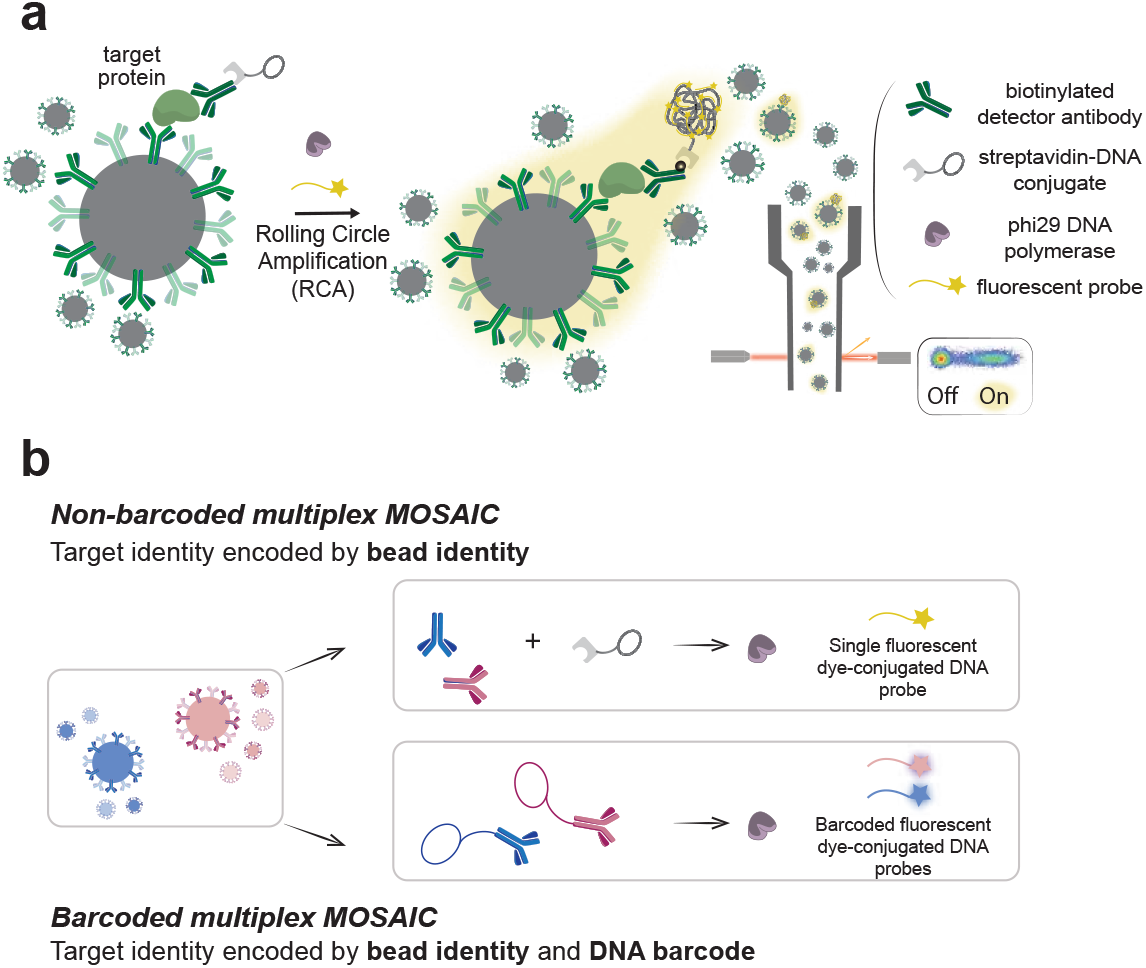
Comparison of MOSAIC and b-MOSAIC. **(a)** MOSAIC workflow. Target proteins are captured on antibody-coated beads, followed by streptavidin–DNA conjugation, RCA, fluorescent probe hybridization, and digital single-color on/off bead readout. **(b)** Multiplexing strategies. In non-barcoded MOSAIC, target identity is encoded by bead identity (e.g., fluorescence color, size, or intensity), while detection is performed using a single-color fluorescent DNA probe. In b-MOSAIC, target identity is dual-encoded by bead identity and a unique DNA barcode, enabling sequence-resolved detection and expanded multiplexing.

**Supplemental Figure S2:**
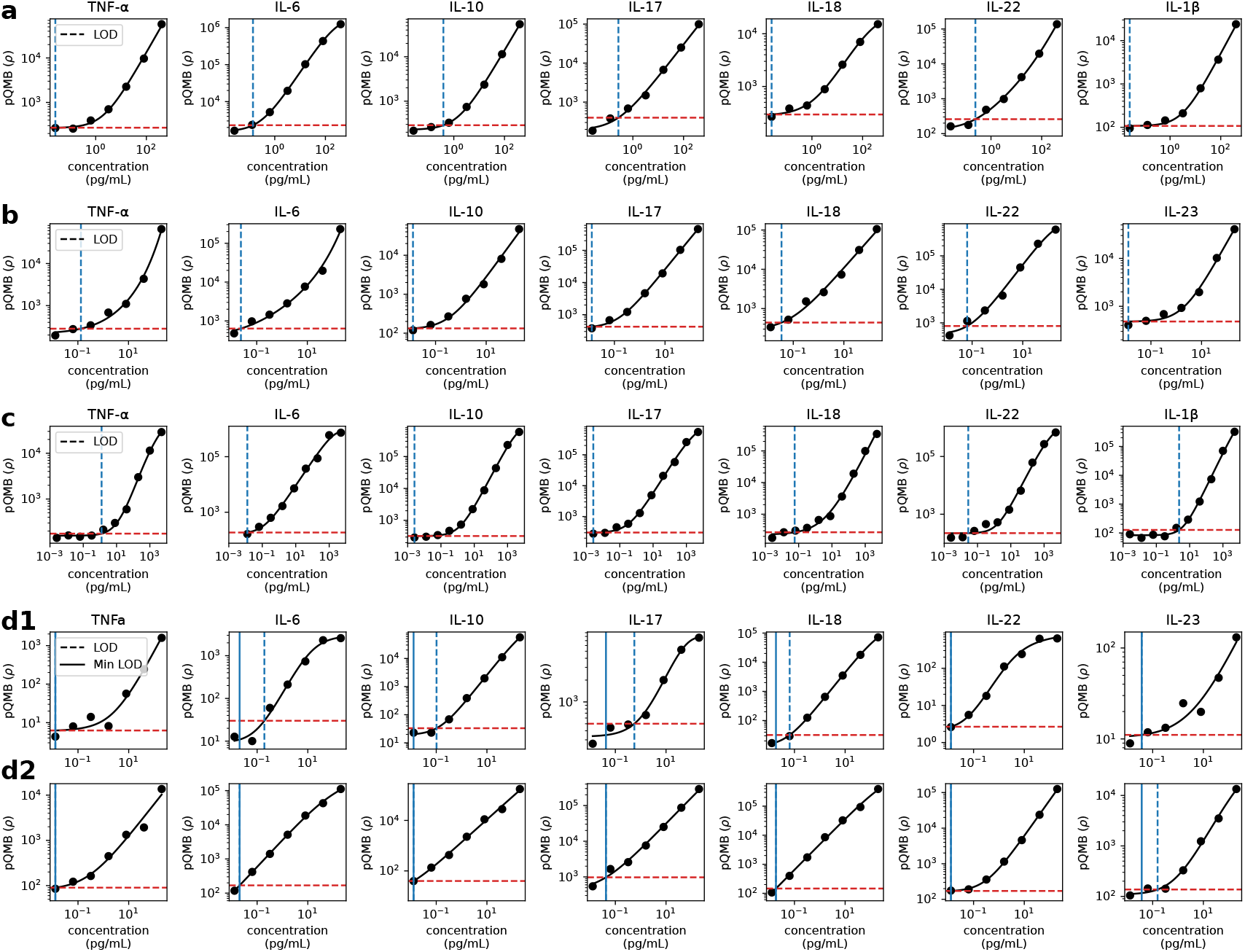
Standard curves used in the study with LOD denoted. **(a)** Standard curves for mouse spike and recovery and limit of detection calculations from **Figure 1e. (b)** Standard curves for human spike and recovery and limit of detection calculations from **Figure 1e** as well as the IBD samples for cohort H1 in **Figure 2b. (c)** Standard curves for mouse cohort M1 data shown in **Figure 2a. (d)** Standard curves for ICI samples in cohort H2 broken down by batch with batches 1 and 2 in panels d1 and d2 respectively.

**Supplemental Figure S3:**
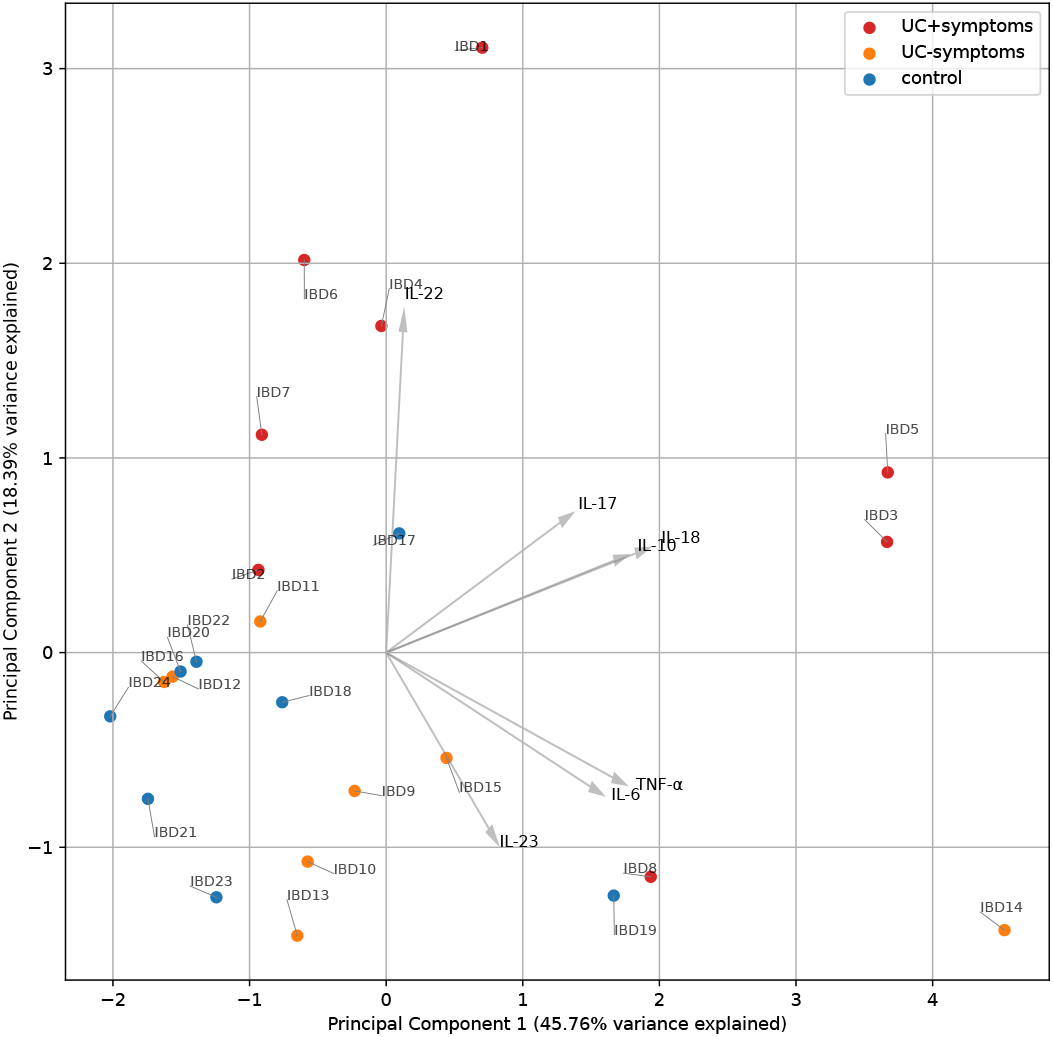
PCA of IBD samples (Figure 2b2) with sample ID annotations.

**Supplemental Figure S4:**
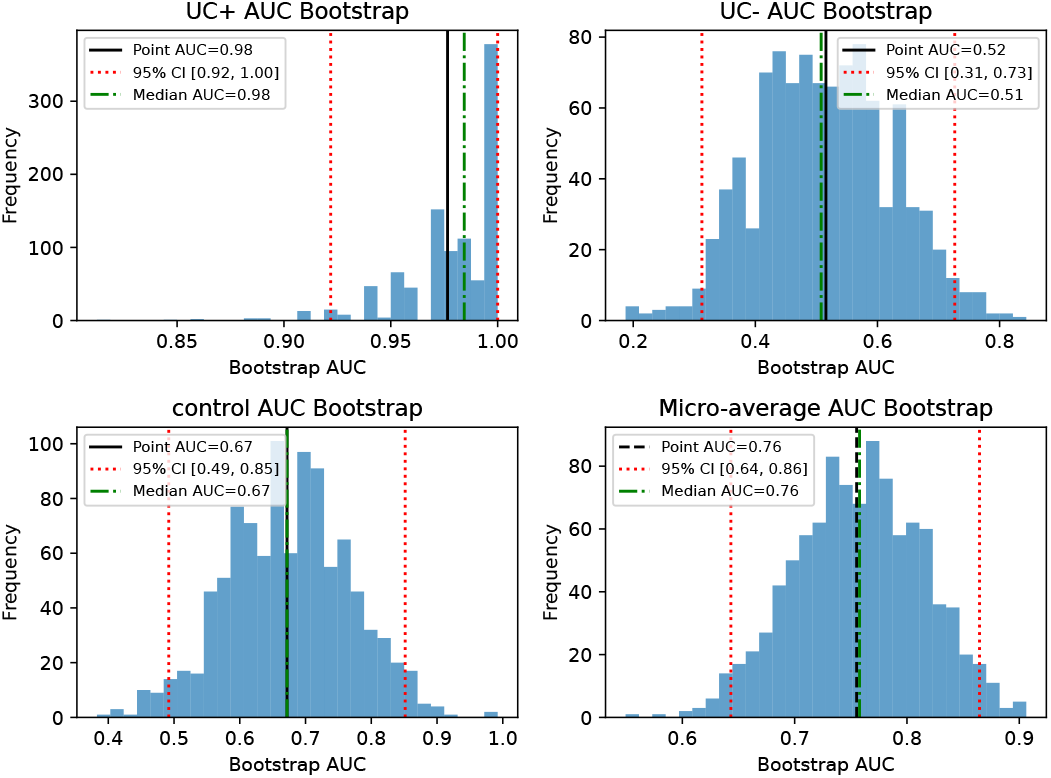
Histogram of bootstrap samples for Figure 2b3.

**Supplemental Figure S5:**
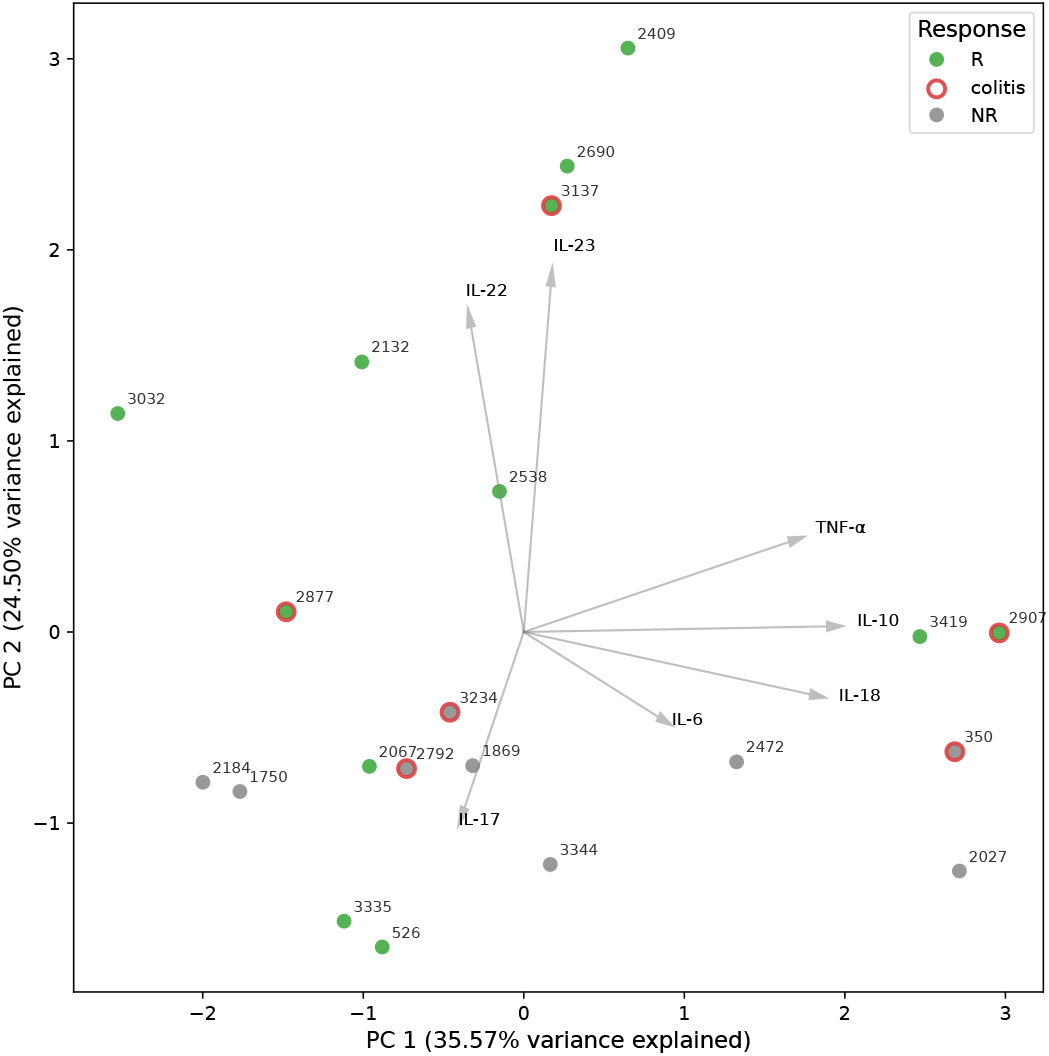
PCA of ICI samples (Figure 2c2) with sample ID annotations.

**Supplemental Figure S6:**
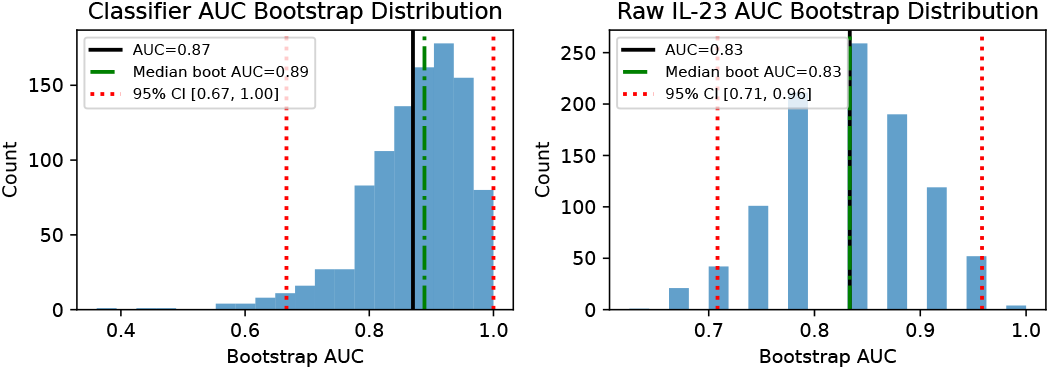
Histogram of bootstrap samples for Figure 2c3.

## Supplementary Tables

**Supplemental Table S1.** Clinical characteristics and medication exposure of the IBD cohort. The cohort included patients with ulcerative colitis with UC+, UC−, and non-IBD controls. Values represent n (%) unless otherwise indicated. Age is shown as median (range). Medication exposure is shown for prior medications documented in clinical history and medications documented at the time of stool collection. Abbreviations: Anti-TNF, anti– tumor necrosis factor; 5-ASA, 5-aminosalicylic acid; PPI, proton pump inhibitor. *5-ASA at collection includes 5-aminosalicylic acid agents explicitly documented as active medications at the time of stool collection. ^#^PPI at collection includes proton pump inhibitors explicitly documented as administered or prescribed at the time of stool collection; historical mentions without confirmation of active use were not counted. Individual patient-level metadata are provided in the cohort metadata tables (Cohort Metadata Excel file, **IBD_cohort** tab).

| Group | N | Sex (F/M) | Age, median (range) | Flare status | Anti-TNF | Thiopurine | Systemic steroid | 5-ASA | Calcineurin inhibitor | Antibiotic | PPI | Probiotic | Allopurinol | Surgery/colectomy |
| --- | --- | --- | --- | --- | --- | --- | --- | --- | --- | --- | --- | --- | --- | --- |
| UC+ | 8 | 2 / 6 | 15 (10–20) | Active flare | Prior medications |  |  |  |  |  |  |  |  |  |
|  |  |  |  |  | 2 (25%) | 4 (50%) | 7 (88%) | 4 (50%) | 2 (25%) | 2 (25%) | 3 (38%) | 0 | 0 | 1 (13%) |
|  |  |  |  |  | Medications at collection |  |  |  |  |  |  |  |  |  |
|  |  |  |  |  | 0 | 4 (50%) | 6 (75%) | 3 (38%)* | 2 (25%) | 1 (13%) | 2 (25%) | 0 | 0 | 0 |
| UC– | 8 | 6 / 2 | 12 (8–17) | No active symptoms | Prior medications |  |  |  |  |  |  |  |  |  |
|  |  |  |  |  | 2 (25%) | 3 (38%) | 3 (38%) | 3 (38%) | 0 | 1 (13%) | 2 (25%) | 3 (38%) | 1 (13%) | 1 (13%) |
|  |  |  |  |  | Medications at collection |  |  |  |  |  |  |  |  |  |
|  |  |  |  |  | 2 (25%) | 3 (38%) | 2 (25%) | 2 (25%) | 0 | 1 (13%) | 0 | 3 (38%) | 1 (13%) | 1 (13%) |
| Control | 8 | 6 / 2 | 10 (3–14) | NA | Prior medications |  |  |  |  |  |  |  |  |  |
|  |  |  |  |  | 0 | 0 | 0 | 0 | 0 | 0 | 0 | 0 | 0 | 0 |
|  |  |  |  |  | Medications at collection |  |  |  |  |  |  |  |  |  |
|  |  |  |  |  | 0 | 0 | 0 | 0 | 0 | 1 (13%) | 1 (13%)# | 0 | 0 | 0 |

**Supplemental Table S2.** Clinical characteristics of the melanoma ICI cohort. Patients were classified as responders or non-responders based on clinical response at 6 months. Values represent n (%) unless otherwise indicated. Stool weight is reported as median (range). irAEs, immune-related adverse events; ipi/nivo, ipilimumab plus nivolumab; nivo/rela, nivolumab plus relatlimab. Neoadjuvant regimens were administered prior to surgical resection. The values shown here represent aggregated summary statistics for the cohort; individual patient-level metadata are provided in the cohort metadata tables (Cohort Metadata excel file, **IT_cohort** tab).

| Group | N | Median stool weight (range), mg | Cutaneous melanoma | Metastatic cutaneous | Patients with irAEs | Pembrolizumab | Ipi/nivo | Nivo/rela | Neoadjuvant regimens |
| --- | --- | --- | --- | --- | --- | --- | --- | --- | --- |
| Responders | 12 | 101.9 (44–140) | 7 (58%) | 5 (42%) | 12 (100%) | 5 (42%) | 4 (33%) | 1 (8%) | 6 (50%) |
| Non-responders | 9 | 130 (44–146) | 4 (44%) | 5 (56%) | 4 (44%) | 1 (11%) | 2 (22%) | 3 (33%) | 4 (44%) |

## Notes

https://github.com/gibsonlab/DIGEST

### Summary of Updates

Hyperlinks to GitHub repo updated to correct address

